# HRDPath: An Explainable Multi-Model Deep Learning Architecture for Predicting Homologous Recombination Deficiency from Histopathology Images

**DOI:** 10.1101/2025.09.24.678258

**Authors:** Chun-I Wu, Kalyan Banda, Rupali Arora, T. Rinda Soong, Marc Radke, Melanie Dillon, Serena Nik-Zainal, Elizabeth M. Swisher, Heba Z. Sailem

## Abstract

Homologous recombination deficiency (HRD) is a critical biomarker for guiding treatment decisions in high-grade serous tubo-ovarian carcinoma (HGSOC), a cancer with few reliable biomarkers. However, existing genomic-based tests for HRD are variable, expensive, and time-consuming. To this end, we developed HRDPath, a novel patient-level deep learning architecture that combines the strengths of two complementary models with a multi-task design, to predict genomically derived HRD status from whole slide images in HGSOC. HRDPath was comprehensively validated across three datasets and benchmarked against leading deep learning models. It achieved an AUC of 0.846, surpassing previously reported H&E-based HRD prediction results for HGSOC images by 0.09, and for the first time, reporting a specificity of 0.938, where accuracy significantly increased when multiple slides per patient were used. Our proposed patient-level approach and interpretability pipeline enhance model trustworthiness and reveal important clinical and biological insights into HRD-positive cancers, highlighting the associated morphological and pathological changes at the cellular and tissue levels. HRDPath is a potentially accessible and scalable digital biomarker that could improve ovarian cancer diagnosis and therapy selection.

## INTRODUCTION

High-grade serous tubo-ovarian carcinoma (HGSOC) is a highly aggressive malignancy with a paucity of available biomarkers for therapeutic intervention. HGSOC exhibiting homologous recombination deficiency (HRD) demonstrates marked sensitivity to targeted treatments such as platinum-based chemotherapy and Poly ADP-Ribose polymerase (PARP) inhibitors^1–3^. Specifically, PARP inhibitors demonstrate synthetic lethality in HRD cancers by inhibiting the repair of damaged DNA, significantly extending progression-free survival and improving long-term outcomes in patients with HRD ovarian cancer^4–6^. Given the significant relationship between HRD status and patient treatment outcomes, the identification of HRD is crucial for precision therapy.

Existing HRD tests analyze both mutations in DNA repair genes, such as *BRCA1/2*, and genomic instability. Examples include: myChoice^7^, which uses *BRCA1/2* alterations and a combined genomic instability score^8,9^; FoundationOne^10^, which assesses *BRCA1/2* alterations alongside loss of heterozygosity and other genomic changes; and HRDetect, a widely accepted tool for HRD detection that has not been adopted clinically due to its dependence on costly whole-genome sequencing^11^. However, current HRD diagnostic tests, such as myChoice, predict a much less robust response to PARP inhibitors in *BRCA1/2* wildtype cancers compared to germline or somatic *BRCA1/2* mutations^4,6,9,12^. Moreover, these tests may be less accessible in regions of the world with limited resources due to their high cost and reliance on advanced genomic technologies.

We hypothesized that using deep learning to predict HRD from hematoxylin and eosin-stained (H&E) WSIs has the potential to become a widely accessible biomarker detection tool, as H&E staining is routinely used in clinical practice worldwide and offers a cost-effective cancer profiling approach. Indeed, previous studies have shown progress in predicting *BRCA1/2* mutations and HRD status from H&E images in breast cancer^21^ but showed limited performance in ovarian cancer with no reported data on method specificity or sensitivity^22^. Additionally, the application of modern slide-based weakly supervised learning approaches where the model takes the entire whole-slide image (WSI) as input and predicts clinically relevant labels such as grade, sub-types or genetic phenotypes are still lacking in the context of HRD prediction in HGSOC^17–19^. Therefore, a robust deep learning model with clinically relevant performance, good explainability, and generalizability has yet to be developed for HGOSC. Moreover, how HRD influences tumor architecture and heterogeneity remains underexplored.

Here we developed a multi-model transformer-based method, HRDPath, for cost-effective HRD detection from gigabyte-sized WSIs of primary or metastatic HGSOC tumors. We utilized a weakly supervised learning approach to identify image regions in WSI that can consistently distinguish patterns and image regions associated with HRD-positive (HRD+) from HRD-negative (HRD-) cancers.

A key feature of HRDPath is its patient-level analysis, aggregating data from multiple slides, without requiring precise anatomical or neoplastic region localization. HRDPath achieved up to 0.846 AUC with 0.692 sensitivity and 0.938 specificity in detecting HRD in an unseen test dataset and achieved comparable performance on two publicly available datasets. To better understand HRDPath performance and the pathological manifestation of HRD in HGSOC and support trust in model predictions, we developed explainability approaches to assess regions that received high attention from the model based on deep-learning features, cancer cell morphology, and tumor composition^10–13^. These analyses consistently revealed that HRD+ HGSOC have distinct morphologies, with polymorphic nuclei which were also confirmed based on the attention of the model. Finally, we demonstrate association between HRDPath predictions and patient outcomes. Our work demonstrates state-of-the-art performance in ovarian cancer and provides significant insights into the pathological changes associated with HRD.

## RESULTS

### Training and evaluation datasets

We utilized an in-house dataset, UWOV, with 152 patients (145 HGSOC and 7 other ovarian cancer patients) as well as two publicly available datasets: TCGA-OV (81 HGSOC patients) and PTRC (93 HGSOC patients) (Figure 1a, and Methods). For our model development, we focused on the UWOV cohort due to its high-quality, large sample size, and the comprehensive information from multiple slides of different anatomical locations per patient, and used TCGA-OV and PTRC as independent validation sets where we only used diagnostic slides from TCGA. The UWOV dataset consists of 720 diagnostic whole tissue section scanned slides from primary debulking resections of HGSOC. The specimens encompassed fallopian tube or ovary (tubo-ovarian: n=480) and extra-ovarian anatomic locations: omentum (n=151), abdomen (n=29), Other Pelvic (n=42), and other locations (n=18) (Figure 1b). HRDetect was used to generate ground truth HRD status for the UWOV dataset^26^ (Methods). 70 patients were classified as HRD-negative (HRD-), while 82 were classified as HRD-positive (HRD+). These cancers were then assessed for HRD-related genomic alterations including *BRCA1* (32.9%), *BRCA*2 (25.7%), and *RAD51C* (2%) mutations (Figure 1c and Methods). Methylation alterations were also assessed, where among the HRD+ cancers, neoplasms from 65 patients (79.3%) harbored either *BRCA1/2*-associated mutations o*r BRCA1* methylation (Figure 1d). we adopted a patient-level training framework, where multiple WSIs from each patient were combined into a single input (Figure 1e and Methods). In the training stage, we considered only patients with diagnostic slides from the primary fallopian tube or ovary (tubo-ovarian), or omentum, as it is the most common metastatic site (146 patients with 631 slide images). The number of slides per patient is comparable between HRD+ and HRD-groups (Overall average:4.3 ± 2.19; HRD-: 4.99 ± 2.07 ; HRD+: 4.52 ± 2.12 ) (Figure 1e). To ensure that HRDPath learns features associated with HRD, rather than patient-specific patterns, we ensured that all slides from a single patient were assigned exclusively to either the training or testing set. Specifically, we used data from 117 patients (80%) with 494 slides (tubo-ovarian: 378; omentum: 116) for training, and 29 patients (20%) with 138 slides (tubo-ovarian: 103; omentum: 35) for validating the model (Supplementary Table 1).

**Figure 1.**
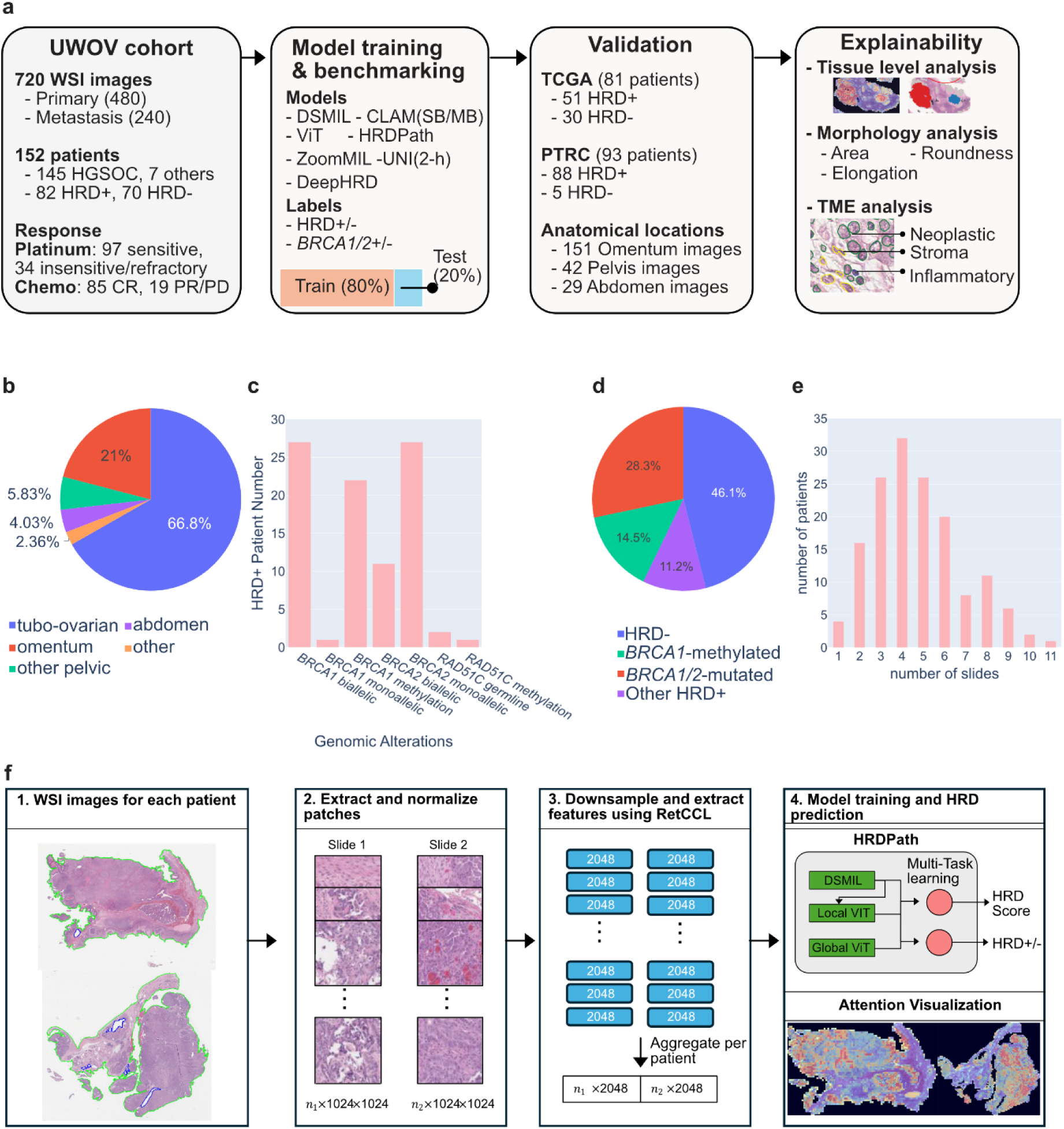
Overview of the HRDPath workflow and study design. **a.** Schematic of the experimental design and dataset composition. Abbreviations: HRD, homologous recombination deficiency; HGSOC, high-grade serous ovarian cancer; WSI, whole-slide image; CR, complete response; PR/PD, partial response/partial denial; DSMIL, Deep Set Multiple Instance Learning; CLAM, Clustering-constrained Attention Multiple Instance Learning; Vil, Vision Transformer; RetCCL, Retrieval with Clustering-guided Contrastive Learning; TME, Tumor microenvironment. b. Distribution of slides by anatomical location. **c.** Genomic stratification of HRD+ patients, including BRCA1/2 mutations, BRCA1 methylation, and RAD51C alterations. d. Proportional distribution of BRCA1/2-associated alterations in HRD+ cases. **e.** Histogram showing the number of slides per patient in the UWOV cohort. **f.** HRDPath pipeline, including patch extraction, feature embedding via RetCCL, model training, and prediction (n;: number of cleaned patches for slide index:i).

Given the large size of WSI images, we extracted smaller patches that were then converted into feature vectors using the CLAM^27^ processing pipeline. We also performed color normalization to ensure transferability across datasets. This pipeline enables the efficient handling of gigapixel images into a series of deep learning-based feature vectors that are then used for the subsequent classification of HRD status.

### HRDPath Development

To predict HRD status from WSIs, we developed HRDPath, a novel weakly supervised learning approach. HRDPath is an end-to-end model that overcomes limitations of previous approaches as it operates without the need for tumor segmentation and has been validated on two independent benchmarking unseen datasets (Table 1-2 and Supplementary Note 1). HRDPath introduces several key novelties, including multi-task learning for predicting binary HRD status as well as continuous HRD score, resulting in more comprehensive evaluations. Additionally, it introduces patient-level learning that increases confidence in prediction by aggregating features from multiple WSIs (Extended Data Figure 1). Moreover, HRDPath adopts an ensemble approach that combines multiple models with different learning strategies employed by Vision Transformers (ViT) ^28^ and Dual Stream Multiple Instance Learning ^29^ (DSMIL models) (Methods). Specifically, HRDPath first utilizes DSMIL, which identifies and aggregates information from key image patches and analyzes how they relate to the remaining patches. Second, to capture local and global information, HRDPath utilizes a ViT model that use and state-of-the-art performance in image analysis.

**Table 1.** Comparison of HRDPath against other existing studies on image-based HRD prediction in Ovarain Cancer (OC). NP (Not performed/provided)

| Study/Method | Loeffler et al.<sup>23</sup> | Zhang et al.<sup>25</sup> | Bergstrom et al.<sup>24</sup><br>(DeepHRD) | HRDPath (Ours) |
| --- | --- | --- | --- | --- |
| **Cancer types considered** | Breast (main focus)<br>OC (train and validate)<br>And 7 others | OC | Breast (main focus)<br>OC (validation only) | OC |
| **Performance on OC (training dataset)** | AUC: 0.63<br>Specificity: NP<br>Sensitivity: NP | AUC: 0.769<br>Specificity: NP<br>Sensitivity: NP | Breast cancer AUC: 0.77–0.85<br>HGSOC: NP | AUC: 0.846<br>Specificity: 0.938<br>Sensitivity :0.692 |
| **Performance on OC (independent dataset)** | NP | NP | NP | Few-shot on PTRC & TCGAOV |
| **Number of OC WSIs** | TCGA (n=520 diagnostic and non diagnostic WSIs) | MSKCC (n=205), Zhongda (n=22) | TCGA subset (n=459 diagnostic and non diagnostic WSIs) + MSKCC (n=141 WSIs) | UWOV (n=720) PTRC (n=198) TCGAOV (n=81 diagnostic) |
| **Model details** | Slide-level attention-based MIL | Two-stages:<br>1) U-Net++ segmentation<br>2) random forest classifier) | Patch-level Ensemble ResNet18 | End-to-end deep learning<br>Multi-task<br>Patient-level |
| **Prediction of continuous HRD score** | No | No | No | Yes (average correlation of 0.69) |
| **Multi-slide aggregation** | No | No | No | Yes (Transformer-based) |
| **Assessment of intra-patient consistency** | NP | NP | NP | Yes |
| **HRD scores considered** | myChoice | myChoice | myChoice or (GIS>40 or *BRCA1/2-mutated*) | - HRDetect<br>- myChoice |
| **Metastases investigation** | No | No | No | Yes (Omentum, abdomen, Other Pelvic sides) |
| **Explainability** | Partially (showing highlighted tumor regions) | Partially (showing highlighted tumor regions) | Partially (showing highlighted tumor regions) | Yes:<br>Neoplastic cell morphology,<br>Tumor microenvironment<br>Clustering of model attention |
| **Limitations** | - Requires large training datasets; - Low ovarian performance.<br>- Limited interpretability | Not end-to-end,<br>- Need segmentation of tumor region;<br>- No direct inference validation | - Patch-level classification<br>- Only select top 5 patches and average their features across an ensemble of models<br>- indirect ovarian validation based on correlation with survival | Main evaluation on HRDetect |

### HRDPath can accurately predict HRD status from histopathological images of HGSOC

When using the DSMIL or ViT model alone for predicting HRD, DSMIL performed better compared to ViT with an AUC of 0.81 with 0.875 specificity and 0.538 sensitivity (Figure 2a and Table 1). This motivated us to develop HRDPath that integrates both models resulting in significant improvement in performance on the test dataset (AUC of 0.846, specificity of 0.938, and sensitivity of 0.692, Figure 1a and 2a). Incorporating a multi-task learning objective proved essential for achieving superior specificity with an absolute improvement of 0.25 and a higher overall performance, reflected by a 0.09 gain in AUC. However, this enhancement came at the cost of reduced sensitivity (Table 2). The performance of HRDPath was generally robust to the proportion of neoplastic cells and cellularity (Extended Data Figure 2a). This suggests that genomic alterations leading to HRD are associated with consistent changes in tissue architecture and visual patterns detectable from H&E images.

**Figure 2.**
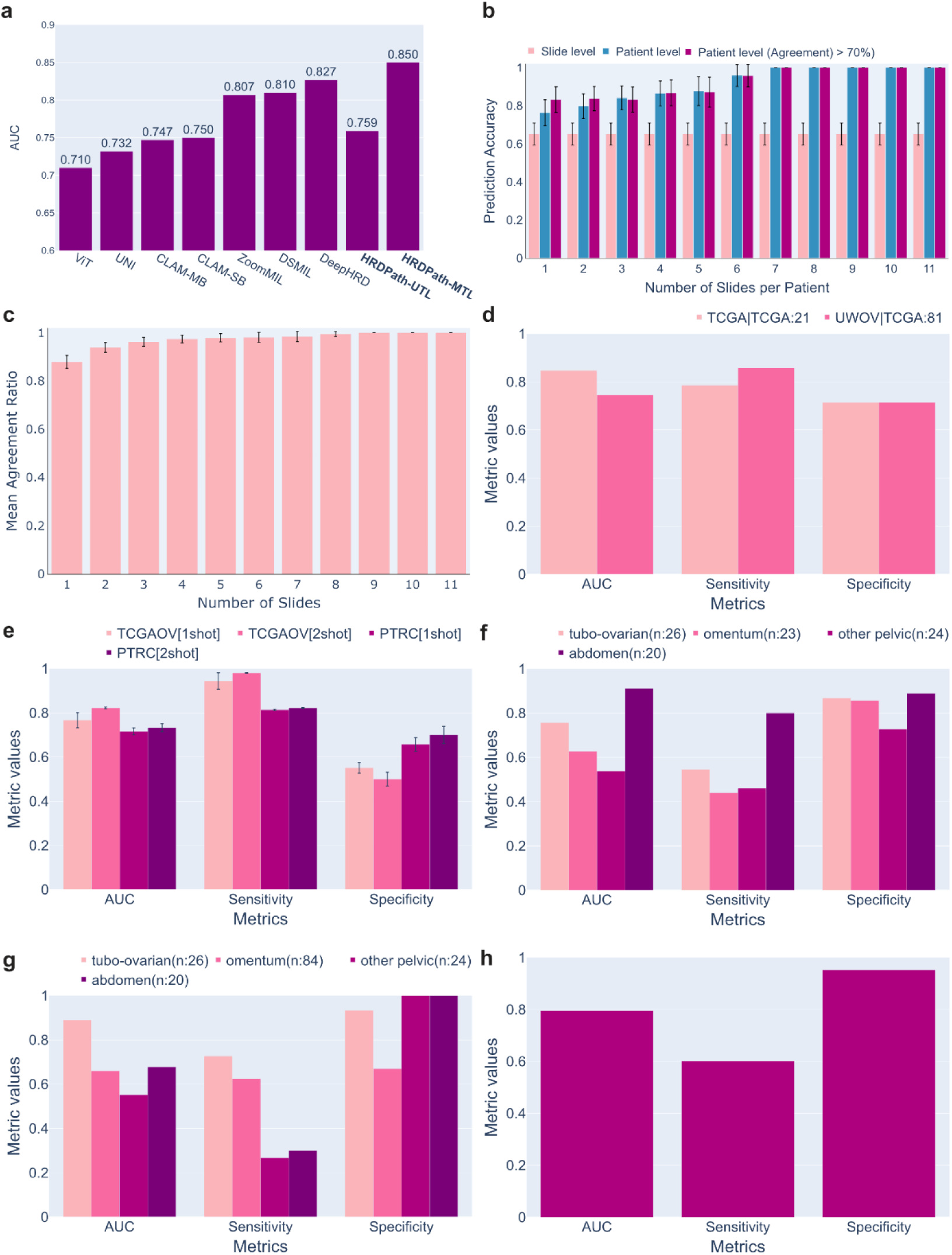
HRDPath accurately predicts HRD on different datasets and anatomical locations. **a.** UWOV Benchmark results from 29 testing patients **b.** Prediction accuracy across 1-11 number of slides as input to HRDPath, error bar indicates .95 confidence interval. **c.** Mean agreement across 1-11 number of slides as input to HRDPath, error bar indicates .95 confidence interval. **d.** HRDPath generalizability experimental results on 21 testing patients from TCGA-OV **e.** Validation of HRDPath using few-shot learning on the TCGA-OV and PTRC-HGSOC datasets. 1-shot and 2-shot indicates that each class is provided with 1 and 2 training cases respectively. **f.** HRDPath (Pretrained on both tuba-ovarian and omental slides) performance across different anatomical locations in the test dataset (29 patients). **g.** HRDPath-primary trained only tuba-ovarian slides can predict HRD status from different anatomical locations. **h.** HRDPath performance on predicting *BRCA*1/2-associated alterations demonstrating its transferability to related classification problems (31 patients) TP: True Positive; FP: False Positive; TN: True Negative; FN: False Negative. UTL: Uni-Task Learning; MTL: Multi-Task Learning.

**Table 2:** HRDPath performance and benchmarking.

| Model | AUC | Sensitivity | Specificity |
| --- | --- | --- | --- |
| ViT | 0.710 | 0.385 | 0.813 |
| UNI | 0.732 | 0.750 | 0.571 |
| CLAM-MB | 0.747 | 0.700 | 0.764 |
| CLAM-SB | 0.750 | 0.615 | 0.875 |
| ZoomMIL | 0.807 | 0.615 | 0.813 |
| DSMIL | 0.810 | 0.538 | 0.875 |
| DeepHRD (k=5, n_ensemble=3) | 0.827 | 0.769 | 0.750 |
| **HRDPath-Uni-task (ours)** | 0.759 | **0.846** | 0.688 |
| **HRDPath-Multi-task (ours)** | **0.846** | 0.692 | **0.938** |

### Evaluation of HRDPath consistency and confidence

As WSIs from multiple tumor locations are often acquired in HGSOC, we sought to evaluate HRDPath consistency when multiple slides are used per patient. We found that HRDPath predictions are generally consistent when using different slides from the same patient, but integrating features from three or more slides can significantly improve model accuracy and consistency (Figure 2b-c and Methods). Importantly, when 6 or more slides were used, accuracy exceeded 95% (Figure 2b). This approach can also be used to derive a confidence score for model predictions by passing individual WSIs through the model and considering the prediction reliable only when at least 70% of slides yield consistent results. These findings underscore the utility of our patient-level approach in increasing the trustworthiness of model prediction.

### Evaluation of HRDPath Against Other Deep Learning Approaches

To rigorously evaluate HRDPath, we benchmarked its performance against widely used and recent deep learning models in histopathology using the same patient cohort. To ensure a fair comparison, all models were trained and tested using the same training/testing split and our proposed multi-task and patient-level learning framework.

Our evaluation included HRD-specific models and state-of-the-art methods with different learning strategies. These methods include the patch-based DeepHRD^24^, multi-scale ZoomMIL^31^ and feature extractions using UNI foundation model^32^. We also included the baseline models of HRDPath: Vision Transformers (ViT)^26^, and DSMIL^27^. We also considered the widely used CLustering-constrained-Attention Multiple-instance learning (CLAM-SB and CLAM-MB), which are both popular multiple instance learning approaches specifically designed for whole slide image analysis. HRDPath outperformed the ResNet-based DeepHRD, particularly in specificity based on our UWOV by 0.19 even when both models were trained using our multi-task, patient-level strategy (Table 2; Figure 2a). It also outperformed the widely used CLAM-SB and its variant CLAM-MB models in predicting HRD with absolute improvements of 0.10 in AUC, 0.063 in specificity, and 0.154 in sensitivity for CLAM-SB (Table 2). Moreover, HRDPath outperformed ZoomMIL, a hierarchical model that integrates features extracted at different image resolutions. Surprisingly, we observe that using the foundation model UNI as a feature extractor results in significantly lower performance (Table 2). These results show that HRDPath consistently outperforms other approaches, supporting its robustness and potential clinical applicability.

### Generalizability of HRDPath

We validated HRDPath on publicly available HGSOC data from TCGA and PTRC. In both TCGA and PTRC, HRD was defined based on myChoice method^33^ where neoplasms were classified as HRD+ if the sum of LOH, LST, and TAI was greater than 42 (Methods). Unlike UWOV, TCGA data comprises 81 diagnostic images from a primary site where only one image is available for each patient (61 HRD+ and 20 HRD-patients)^34^. PTRC-HGSOC comprises 198 slides (106 tubo-ovary and 92 omentum) from 93 patients from different hospitals with highly imbalanced HRD distribution (88 HRD+ and 5 HRD-patients). Despite the small size of the TCGA dataset, HRDPath achieved comparable performance in predicting HRD status when trained on TCGA, consistently outperforming other models, achieving 0.847, 0.714, and 0.786 in AUC, specificity, and sensitivity, respectively (Extended Data Figure 2d; Supplementary Table 3). These results confirm that HRD status defined based on different HRD assays can be robustly predicted from H&E images.

To further confirm HRDPath generalizability, we evaluated its performance on independent cohorts using few-shot learning, images of metastatic neoplasms, and classification of *BRCA1/2* alterations. We employed 3-fold few-shot learning where data from only one or two support cases (i.e. patients) per class to determine if HRDPath can be applied without significant finetuning. HRDPath, initially trained on the UWOV dataset, accurately predicted HRD in the TCGA and PTRC-HGSOC datasets after being fine-tuned with 1-3 example images with AUC reaching 0.82 and specificity reaching 0.98 (Figure 2e; Supplementary Table 4). We found that color normalization significantly improved HRDPath generalizability across different datasets (Extended Data Figure 2c; Supplementary Table 5). HRDPath showed strong performance in predicting HRD from abdominal metastatic WSIs despite no training data from this site, though performance was lower on Other Pelvic samples (Figure 2f-g and Supplementary Note 2). It also classified *BRCA1/2* alterations with good accuracy (AUC = 0.8, Figure 2H and Supplementary Note 3). These findings highlight the robustness and generalizability of HRDPath and suggest concordance between HRDetect and myChoice scores. Most importantly, the first demonstration of HRD prediction in HGSOC using few-shot learning with as few as one sample per class, a key step toward clinical translation.

### Characterization of pathological and morphological changes identified by HRDPath

To increase the confidence in the reliability of HRDPath predictions, we analyzed tissue architecture to explain HRDPath predictions. Although HRDPath did not receive any information on neoplastic regions, it successfully identified that visual patterns associated with HRD were generally located in neoplastic regions (average Pearson Correlation between neoplastic proportion and model attention=0.34 and Figure 3a).

**Figure 3.**
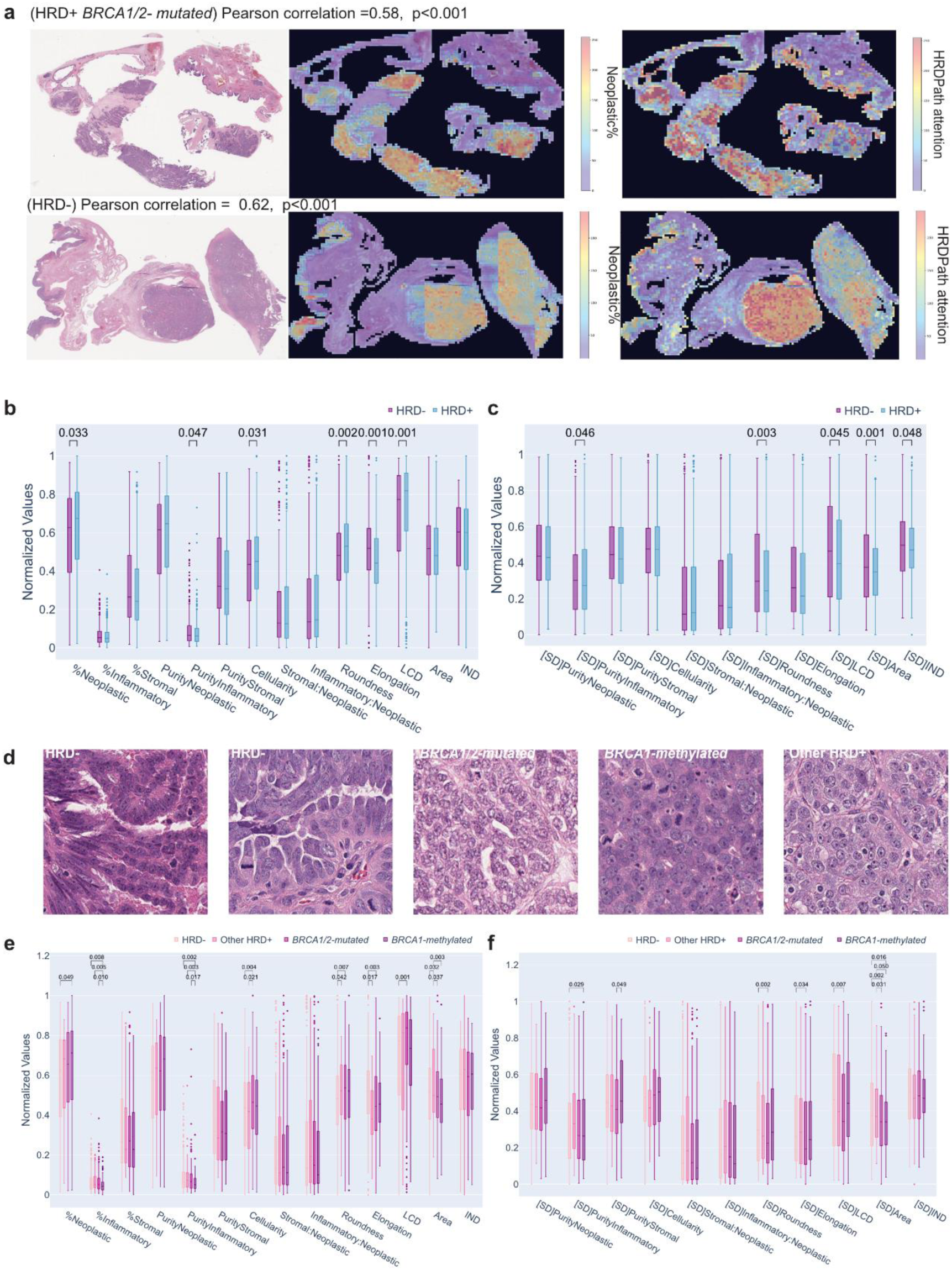
Characterization of morphological differences between HRD status and genomic alterations. **a.** Example WSI images of HRD+/-neoplasms (left), heatmaps of the corresponding neoplastic proportion (middle), and heatmap representing HRDPath attention (right). Pearson correlation is between patch-wise HRDPath attention and Neoplastic%. **b,e.** Distribution of average of nuclear morphology per patient across HRD status (b) and genomic alterations (e). **c,f.** SD of nuclear morphology across HRD status (c), and genomics status (f). **d.** Example regions across genomic alterations. *P-values* are calculated with the two-sided T-test and shown only if *p-value* < 0.05.

Using the HoVer-NeXt^35^ model we performed single-cell morphological analysis on 720 slides, while restricting to regions that received high attention (≥ 0.7) from HRDPath. We considered neoplastic, inflammatory, and stromal cell types as these were the most accurately predicted by the model as evaluated by a pathologist (Methods). We computed various features to quantify tumor composition, including proportion of different cell types, cellularity cell purity, which measures the extent to which a single cell type dominates a specific image region (i.e., patch), and the ratio between different cell types which has been reported to be associated with clinical outcomes in HGSOC^36^. Additionally, we measured the mean and standard deviation (SD) of various morphological features of neoplastic nuclei, including roundness, elongation, area, local cell distance (LCD), and immune-to-neoplastic cells Distance (IND) (Methods).

We observed a significantly higher percentage of neoplastic cells in regions associated with HRD+ HGSOC (p≤0.05; Figure 3b). These differences were not due to variations in the overall neoplastic proportion between HRD+ and HRD-cancers, as no significant difference was observed when considering the entire WSI (Extended Data Figure 2a). Additionally, neoplastic nuclei exhibited significantly lower elongation, higher LCD, and roundness in HRD+ cancers which may indicate loss of epithelial morphology (p≤0.001, p≤0.001, and p≤0.05 respectively; Figure 3b). We also observed a more uniform morphology as indicated by the lower SD of roundness, area and inflammatory purity (Figure 3c). These results suggest that genomic alterations linked to HRD are mainly associated with an increased proportion of neoplastic cells and polymorphic nuclear morphology.

As *BRCA1/2*-associated alterations are most common in HRD+ patients, we sought to evaluate whether *BRCA1/2*-associated alterations exhibit differences in morphology. The most consistent changes observed across different HRD+ alterations were increased nuclear roundness and decreased elongation; however, this increase was not significant in the *BRCA1*-methylated group (p≤ 0.05, p≥0.05 (0.07), p≤0.05 respectively; Figure 3d-e). *BRCA1/2*-mutated and other HRD+ HGSOC were associated with an increased proportion of inflammatory cells, which was more significant when only the neoplastic regions were considered (Figure 3e and Extended Data Figure 2e-g). We observed a decrease in the mean and SD of nuclear area in HRD+ cancers with *BRCA1/2* alterations which were most significant in *BRCA1-*methylated cancers (Figure 3e-f). Notably, *BRCA1/2*-mutated cancers had a significantly higher median of LCD compared with HRD-, whereas *BRCA1*-methylated cancers had the lowest LCD (Figure 3e). These results suggest that different genomic alterations underlying HRD can vary in morphology and immune microenvironment but are generally characterized by rounder nuclei and more homogenous morphology.

### HRDPath reveals morphological patterns across primary and metastatic neoplasms

As HRDPath accurately detected HRD+ cancers from images of metastatic HGSOC, we investigated the morphological differences in HRD+/-between primary (tubo-ovarian) and omental sites. The metastatic regions had a higher stromal-to-neoplastic ratio, lower cellularity, higher proportion and purity of inflammatory cells, and lower elongation regardless of HRD status (Figure 4a). As expected, metastatic regions also showed higher variability (SD) in inflammatory purity, stromal to neoplastic cell ratio, and IND (Figure 4b). HRD-omental metastases exhibit a significant increase in roundness and its SD compared to HRD-primary neoplasia (p≤0.05; Figure 4a-b). On the other hand, HRD+ omental metastases exhibited significantly larger neoplastic cells with lower elongation, and lower SD of stromal purity compared with HRD+ primary neoplasia (p≤0.05; Figure 4a-b). These results confirm that HRD alterations are associated with consistent changes in certain morphological features across different disease sites and indicate the unique characteristics of the tumor microenvironment in metastatic HGSOC.

**Figure 4.**
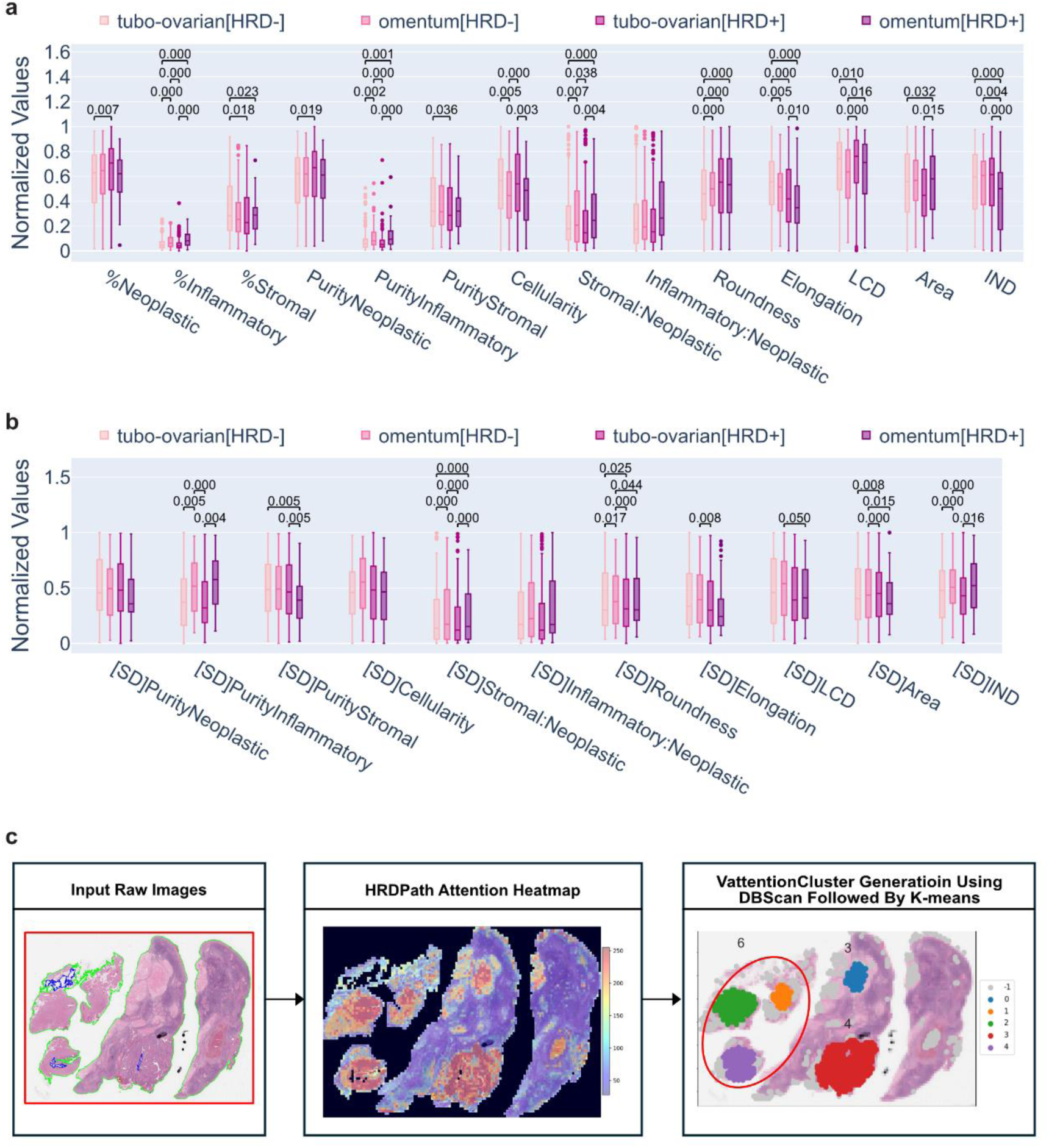
Morphological characterization across primary and metastatic sites and VAttentionClusters. **a.** Nuclear morphology distribution across primary and metastatic sites **b.** Nuclear morphology standard deviation distribution across primary and metastatic sites **c.** VAttentionClusters generation pipeline. **1)** HRDPath attention is first clustered according to spatial distribution using DBScan and outliers are removed, 2) K-Means clustering (k=B) is applied to attention clustered obtained from 1) considering their attention-weighted RetCCL features to form VattentionCluster (Methods). *P-values* are calculated with the two-sided T-test and shown only if *p-value* < 0.05.

### Systematic analysis of tissue-level and tumor microenvironment phenotypes

To gain further insights into tissue-level phenotypes associated with HRD and enhance trust in model predicitons, we developed a novel approach to determine key phenotypes based on deep learning features (Figure 4c). We first identified patches that received high attention from HRDPath (Methods). Unsupervised DBscan clustering was applied to patches considering the spatial arrangement to form attention regions. These regions were then grouped into eight clusters using k-means clustering based on their RetCCL-derived visual features and attention scores, which we refer to as VAttentionClusters. Inspecting these clusters qualitatively and quantitatively revealed that VAttentionClusters 1,2,3, and 4 predominantly consisted of neoplastic cells and exhibited distinct phenotypes (Figure 5 and 6a-b). VAttentionCluster 1 was significantly enriched in HRD-HGSOC whereas VAttentionCluster 3 was significantly enriched in HRD+ cancers (Figure 6c). Nuclei in VAttentionClusters 1 and 4 were significantly less elongated and rounder than VAttentionClusters 2 and 3 (Figure 6c). Moreover, nuclei in VAttentionCluster 3 had the lowest local cell density and were most enriched in tumors harboring *BRCA1/2* alterations (Figure 6d). Interestingly, VAttentionCluster 2, although significantly enriched in Other HRD+ cases, exhibited high nuclear elongation and the lowest roundness among neoplastic clusters (Figure 6b-d). These results suggest that genetic alterations associated with HRD are correlated with distinct tumor morphologies.

**Figure 5.**
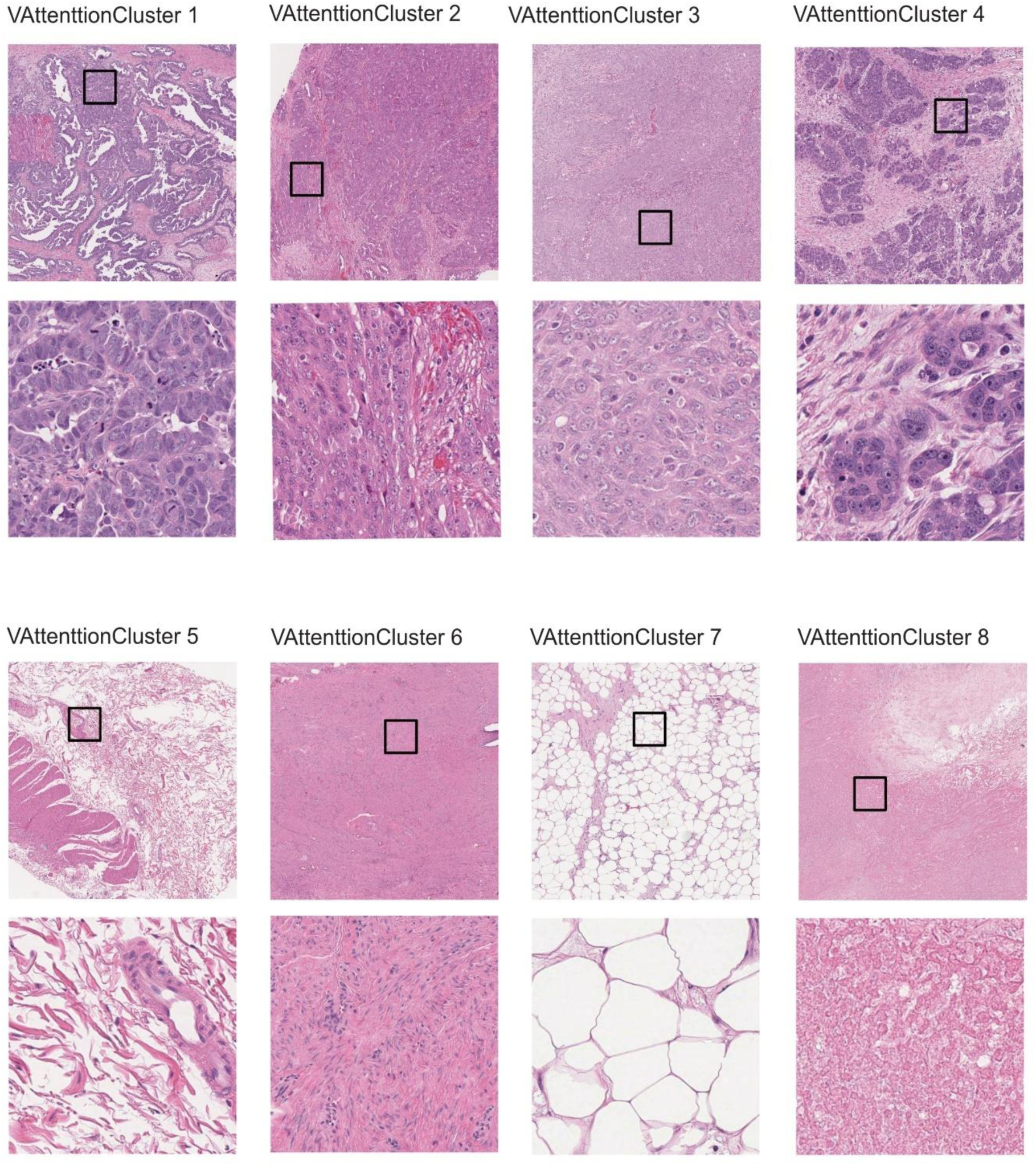
Representative images of different VAttentionClusters. The top row shows a zoomed-out image and the bottom row displays a zoomed-in image of the region outlined by the black box in the top row. VAttentionClusters 1-4 are dominated by neoplastic cells. VAttentionClusters 5-6 is dominated by stromal and fibrotic tissue. VAttentionCluster 7 compromises adipose tissue, while VAttentionCluster 8 compromises infracted tissue.

**Figure 6.**
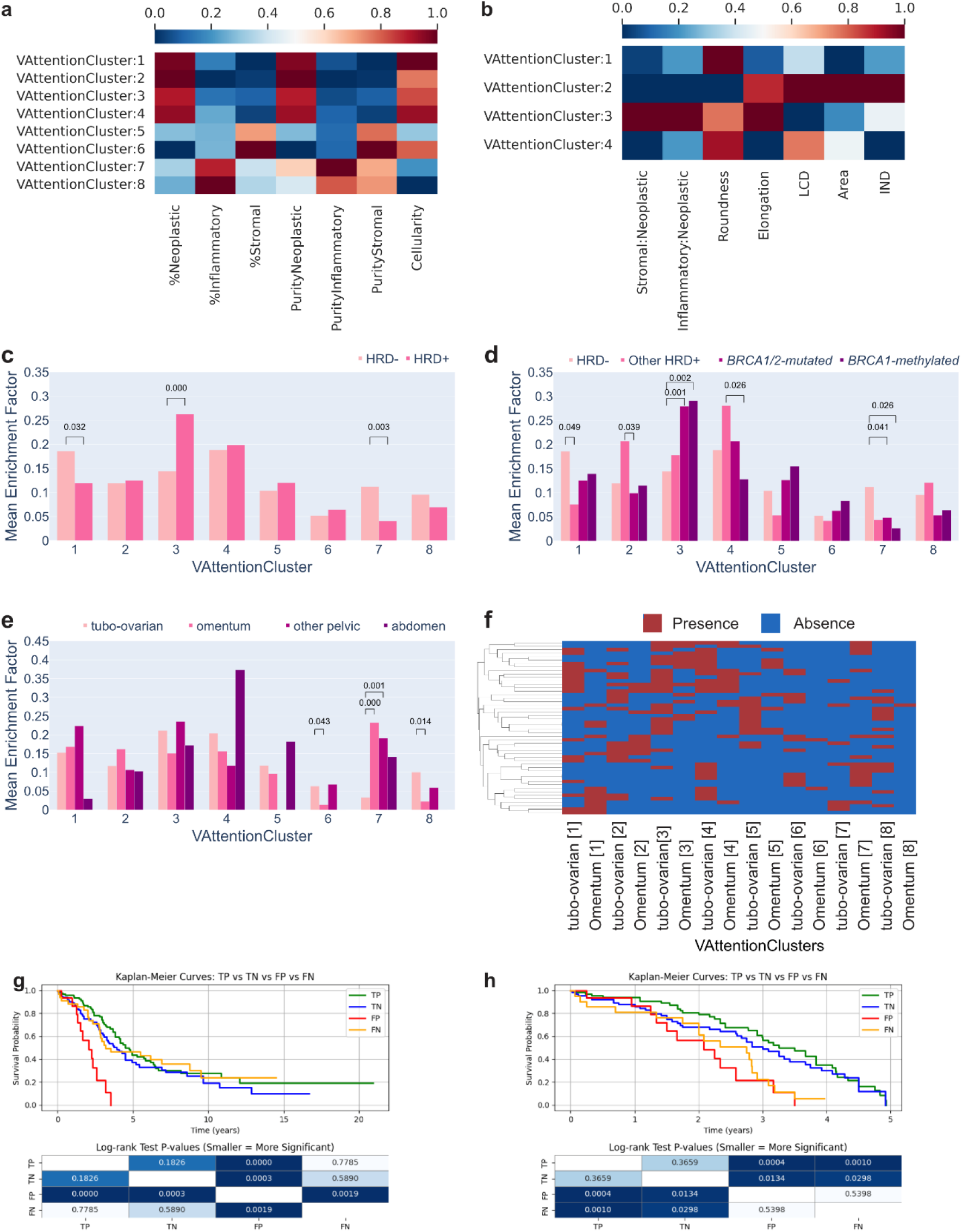
VAttentionClusters enrichment factor distribution across HRD status, genomic alterations, and anatomical locations and response groups. **a.** Cell distribution across VAttentionClusters **b.** Morphological characterization of VAttentionClusters (neoplastic clusters). **c-e.** VAttentionClusters enrichment factor across different HRD status (c), genomic alterations (d), anatomical locations (e) **f.** Enrichment of VAttentionClusters in primary and omental neoplasia where cancers with similar profiles are grouped by hierarchical clustering. *P-values were calculated with the two-sided T-test (c-f). P-values* were calculated with the two-sided T-test. g-h. A Kaplan Meier survival analysis of discordant cases based on overall survival (g) and patients with <5 years survival (h). Shown heatmaps represent log-rank test p-values. TP (True Positives), TN (True Negatives), FP (False Positivies), FN (False Negatives).

Our VAttentionClusters revealed that HRDPath also identified several tumor microenvironment signatures associated with HRD status. Specifically, VAttentionClusters 5-8, were characterized by low neoplastic cell proportions and cellularity but were dominated by necrotic, stromal, fibrotic, or adipose tissue regions (Figure 5). Through qualitative assessment, we found that VAttentionCluster 7 was mainly composed of adipose tissue and was significantly associated with HRD-cancers (Figure 6c). Conversely, VAttentionClusters 5 and 6 were primarily composed of stromal cells with high fibrosis present in VAttentionCluster 5 supported by its lower cellularity and high stromal proportion (Figure 5 and 6a). While these clusters were present in primary sites, VAttentionCluster 5 was also present in the omentum and abdomen while VAttentionCluster 6 was present in Other Pelvic metastases (Figure 6e). VAttentionCluster 8, on the other hand, was characterized by infarcted tissue and fibrosis with the lowest cellularity and was enriched in both HRD- and Other HRD+ cancers (Figure 6a-d). These results suggest that HRD+ cancers display location-specific changes in their tumor microenvironment.

Next, we investigated whether VAttentionClusters tended to localize specific anatomical regions. All neoplastic clusters were present across different locations, except for VAttentionCluster 1, which showed a low enrichment factor in the abdomen (Figure 6e). In contrast, abdominal metastases were mostly enriched for VAttentionCluster 4 (Figure 6e). While the stromal VAttentionCluster 5 and 6 were present in primary sites, Stromal VAttentionCluster 5 was absent in the Other Pelvic metastases but present in the abdominal metastases. On the other hand, Stromal VAttentionCluster 6 was detected in the primary sites and Other Pelvic metastases but absent in the abdominal metastases, suggesting location-specific signatures. Adipose VAttentionCluster 7 had a very low enrichment factor in primary sites, while necrotic VAttentionCluster 8 was mostly enriched in primary or Other Pelvic sites (p≤0.001; Figure 6e). We also observed that if VAttentionCluster 2-4 is present on the primary site, it is likely to be present on the omental site in that patient (Figure 6f). These results suggest that phenotypic changes associated with HRD are shared between primary and metastatic locations, but that some tumor microenvironment phenotypes might be location-specific.

### Association of HRDPath predictions and patient outcomes

We found that differences in genomic alterations and morphological heterogeneity could partially explain discordant cases where image-based HRD prediction did not match the genomic-based HRD status (Supplementary note 4). Therefore, we sought to investigate whether HRDPath could stratify patient outcomes. Interestingly, false positive cases (HRDPath+/HRD-) were associated with significantly worse survival when considering both UWOV and TCGA cohorts (p≤ 0.01, Figure 6g). Moreover, the false negative cases (HRDPath-/HRD+) showed significantly worse survival than the true positives and true negatives when considering patients with less than 5 years of survival only (p≤0.05 and Figure 6h). This suggests that image-based predictions by HRDPath capture clinically meaningful phenotypes not reflected in genomic-based HRD assays, highlighting potential for broader prognostic relevance.

## DISCUSSION

In this study, we developed HRDPath, a novel multi-model architecture to predict HRD status using H&E images of HGSOC. HRDPath marks a significant advance with an AUC of 0.846 and specificity of 0.94. We validate and show that it performs comparably well on unseen images from TCGA-OV and PTRC even when only finetuned on one image per class using few few-shot learning paradigm. We also show for the first time that HRD can be predicted from metastatic tissue slides. HRDPath also achieved state-of-the-art performance in predicting *BRCA1/2* alterations outperforming existing methods^37–40^. Additionally, it is the first to perform patient-level classification by incorporating data from multiple slide images and anatomical locations to improve model performance and we demonstrate that this leads to very high accuracy when 6 or more slides are included. Our comprehensive phenotypic profiling reveals new important insights on morphological correlates with HRD at cellular and tissue levels which can be exploited in therapy development and target discovery.

Although deep learning models showed great promise in other cancer types^41–43^, their application to ovarian cancer has yielded limited performance with reported AUC values ranging from 0.64 to 0.76 and no reports of specificity^23,25,44^ This is likely due to architectural constraints. For instance, DeepHRD utilized a computationally intensive, patch-level classification strategy that not only ignored broader contextual information but also relied on top five patches for its prediction. Furthermore, a shared limitation among these previous methods is the lack of evaluation on truly independent patient cohorts. Another approach required manual tumor region segmentation and reliance on manual feature extraction, which were then fed into a random forest classifier hindering the translation of this approach ^25^. In contrast, our approach only requires patch extraction using CLAM pipeline and feature extraction using RetCCL feature extractor. As such, we anticipate that our approach is more robust to inter-dataset variability as it requires minimal preprocessing. Interestingly, using UNI foundation models as feature extractors, instead of RetCCL, did not improve performance based on our data. Importantly, our evaluation of HRDPath on both unseen data within the same cohort and on independent cohorts further underscores the importance of color normalization in addressing domain shifts, particularly in one-shot or few-shot learning scenarios. Overall, our study addresses key gaps in HRD prediction and offers a computationally efficient, end-to-end framework suitable for broader application.

The integration of ViT and DSMIL into HRDPath architecture allowed effective detection of different pathological changes associated with diverse HRD alterations that include copy number amplifications, mutations, and large genomic structural phenotypes. HRDPath not only extracts meaningful features from gigapixel-sized whole-slide tissue images but also identifies the most relevant patches and regions among thousands. Specifically, DSMIL selects relevant regions, thereby guiding the transformer model’s attention, while the transformer-based feature fusion method to integrate slide-level and patient-level information. We found that this patient-level and multi-model approach can significantly improve the overall performance of our model, where we found model predictions become stable with three or more slides. Interestingly, our patient-level approach also allowed us to quantify the variability in model predictions, showing that even with one slide, HRDPath tends to be consistent in its predictions. Moreover, HRDPath outperformed more recent advanced approaches such as the ZoomMIL model, which identifies relevant patches from low-resolution images and analyzes their features at higher resolution. This may be because HRD-associated alterations are more discernible at the single-cell level. Our findings are also consistent with results from our AI in Histopathology Explorer study, where we observed that multi-model architectures often outperform single-model approaches and are less sensitive to dataset size based on comparison of more than 1,400 published studies^45^. These results suggest that diversifying model learning strategies may be more effective than focusing on building a single, highly specialized models.

Our explainability framework allows the model to highlight relevant regions and phenotypes, providing clinicians with critical insights to evaluate and validate the trustworthiness of individual predictions. Moreover, many of the differences identified by HRDPath were consistent with previous observations that *BRCA1/2*-mutated ovarian cancers have a higher frequency of certain pathological and invasion patterns^47,48^. Importantly, we also found that *BRCA1*-methylated cancers exhibited distinct morphologies. Deep learning models allowed systematic and quantitative discovery of the multitude of phenotypes associated with HRD+ cancers using WSI images, integrating information from thousands of patches in a non-linear manner which might be challenging to detect by the naked eye alone. Our results are generally consistent with the features reported in HRD+ breast cancers increased fibrosis, and polymorphic nuclei^49^. The decrease in the overall proportion of stromal cells in HRD+ HGSOC might reflect a more complex interplay of features, as the reduced stromal cell count could be indicative of increased fibrosis as it was also associated with reduced cellularity. This interpretation is supported by our tissue-level analysis, which showed a higher enrichment of fibrosis-associated VAttentionClusters 5 and 6 in cancers with *BRCA1/2* genomic alterations. Unlike previous reports in HRD+ breast cancers, our analysis does not support increased necrosis as a distinguishing feature of HRD+ HGSOC, as reported in HRD+ breast cancers. Our explainability pipeline can be potentially used to discover novel phenotypic differences but also to increase model trustworthiness, by highlighting regions aligned with model predictions.

Analysing single-cell morphology and neoplastic composition revealed that nuclei are consistently more round and less elongated in HRD+ primary and metastatic cancers. The increased nuclear roundness may be due to higher proliferative activity resulting from DNA repair impairment, as dividing cells often exhibit rounder shapes during mitosis^50^. Moreover, the high proliferation rate, as reflected by the higher neoplastic density, can disrupt epithelial polarity, as well as cell adhesion and cytoskeleton^51^. Interestingly, we identified a subset of HRD+ cancers that showed significantly increased elongation and reduced roundness, especially in cancers with no *BRCA1/2*-associated alterations. This could be associated with epithelial-to-mesenchymal transition which has been linked to differential DNA damage repair response^52,53^. Intriguingly, misclassified cases, and especially HRDPath+/HRD-, were associated with significantly worse patient outcome. Evaluating how morphological patterns can be leveraged to refine prognostic stratification in HGOSC would be an interesting future direction.

The evaluation of our balanced in-house UWOV and TCGA, along with additional evaluation on metastatic slides, provides strong evidence of the generalizability of our approach. While HRDetect provides a well-validated genomic signature with demonstrated correlation to *BRCA1/2* deficiency^54–56^, we recognize that it is not FDA-approved. Nonetheless, the validation on two independent cohorts (TCGA and PTRC) where myChoice labels were available addresses this limitation and demonstrates consistent performance based on few-shot learning. Broader validation across more diverse patient populations and balanced datasets will be necessary to advance HRDPath translation potential, especially given that PTRC dataset included only five HRD-patients. Future work includes additional validation of HRDPath using multiple FDA-approved assays will be important to support clinical translation.

We believe that HRDPath has the potential to be translated into a digital biomarker for detecting HRD in ovarian cancer patients, integrating histopathology images with a transformer-based deep learning model for rapid assessment of genetic signatures. Our approach is the first to demonstrate significant improvement over existing methods and predict HRD from metastatic tumors, thus expanding its potential clinical utility beyond primary sites. Additionally, our morphological analysis uncovered previously unrecognized features of HRD+ HGSOC, offering important insights into cancer evolution and tumor microenvironment interactions. Given that HGSOC remains a poorly understood malignancy with one of the lowest survival rates, these findings are both timely and impactful. We envision that HRDPath can in future translate to a surrogate biomarker before genomic testing or offer cost-effective biomarker in low- and middle-income countries toward improving patient outcomes. Together, these advancements establish HRDPath as a potentially cost-effective tool for refining ovarian cancer diagnostics and treatment strategies.

## METHODS

### Study Design

In this study, we developed HRDPath, a patient-level deep learning model for predicting HRD status from histopathological images of HGSOC (Figure 1a). HRDPath was evaluated on UWOV and a public dataset TCGA-OV, with performance on unseen data benchmarked against CLAM-SB, DSMIL, ViT, DeepHRD, ZoomMIL, and UNI. To assess robustness, the model was tested across tumor sites, including primary and metastatic lesions (omental, abdominal and other pelvic). We validated HRDPath generalizability to new datasets through few-shot learning. Additionally, we trained a specialized model for predicting *BRCA1/2* alterations using transfer learning that can provide further complimentary biomarkers for treatment and targeted selection for clinical trials. Finally, we demonstrated HRDPath’s trustworthiness through multi-scale analysis of the model attention as well as analysis of model prediction consistency when different WSIs were fed for the same patient. This allowed characterization of attention-driven phenotypes of the tumor microenvironment using a clustering pipeline which were correlated with HoVer-NeXt derived cell morphological features through statistical analysis.

### Patient Cohort

#### University of Washington Ovarian (UWOV) dataset and patient cohort

Patients diagnosed with high-grade ovarian cancer, undergoing surgery, and seeking treatment at the University of Washington were enrolled in a biorepository approved by the Institutional Review Board (IRB) (applicable registration). All patients provided written informed consent prior to enrolment. In this study, 212 representative patients with 222 samples were selected. Of these, 152 patients had both HRD scores generated, and H&E images digitized (Supplementary Table 1). All patients were followed until death and, if alive, censored at the last follow-up.

In this dataset, HRD status is defined using HRDetect algorithm. HRDetect is a whole-genome sequencing-based algorithm that predicts cancers with high, intermediate or low HRD score ^58,59^. It is a weighted model that combines the contributions of six distinguishing mutational signatures to produce a signal score predictive of a defect in the homologous recombination DNA repair pathway. Following signature assignment, the contributions of substitution signatures SBS3 and SBS8, rearrangement signatures RefSig R3 and R5, loss of heterozygosity index and proportion of small deletions in regions of microhomology were used to calculate the HRDetect probability score. This score is predictive of the response to PARP inhibitors in HGSOC (paper under second round of revision).

#### TCGA Ovarian cancer whole-slide image dataset

This study was validated on publicly available H&E diagnostic slides and whole genomic sequence from the TCGA Ovarian dataset (https://portal.gdc.cancer.gov/). 81 diagnostic slides were processed using the same pipeline for UWOV (Supplementary Table 1). HRD score was sourced from previous studies^60^.

#### PTRC-HGSOC whole slide image dataset

198 H&E slides from PTRC-HGSOC were processed using the same pipeline. The HRD score was defined as the sum of LOH, LST, TAI provided in the original paper^30^.

### Image Preprocessing

To make WSI images suitable for deep learning models. We adapted the processing steps of CLAM. Each WSI is cut into a sequence of square patches of 1024 pixels at 40x resolution (0.25 µm/pixel) at the selected level dimensions. The level dimension that has a resolution closest to 1e^8^ was selected. Background patches were filtered using median thresholding. To mitigate the effect of H&E staining variation within the dataset, the patches were normalized based on slide-level statistics. For each whole-slide image, the mean and standard deviation of the red, green, and blue channels were computed across all pixels within the slide. Subsequently, each pixel value in the corresponding patch was standardized by subtracting the channel-specific mean and dividing by the channel-specific standard deviation. Each patch was resized via bi-linear interpolation to 384 pixels, which is equivalent to ∼15× magnification and is sufficient to retain nuclear and cytological features needed for cell-level analysis. Then, patches were processed using RetCCL, a self-supervised model trained on 32000 pan-cancer whole slide images ^61^, to obtain 2048 features per slide.

### Patient-Level Multi-task Learning

We included HRD score prediction as a complementary task because the HRD+ group is heterogeneous. Some HRD+ patients exhibit borderline HRD scores that may reflect distinct morphological characteristics. By jointly optimizing for both classification and regression, the model is encouraged to capture finer-grained morphological cues, enhancing generalizability and interpretability. Specifically, our framework simultaneously performs (i) HRD status classification, a binary prediction task (HRD+ vs HRD−), and (ii) HRD score regression, a continuous prediction of the homologous recombination deficiency (HRD) score derived from genomic data, with both tasks given equal weights using the following objective function:

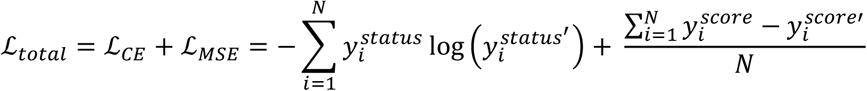

Where ℒ_𝑡𝑜𝑡𝑎𝑙_ represents the combined multi-task loss, consisting of classification cross entropy loss ℒ_𝐶𝐸_ and regression mean-squared error loss ℒ_𝑀𝑆𝐸_. _Here,_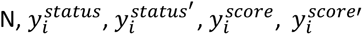 _denotes_ the total number of training samples, HRDPath HRD status prediction, HRD status ground truth, HRDPath score prediction, HRD score ground truth respectively.

### HRDPath Architecture

HRDPath integrates DSMIL and Vision Transformer with proposed local window attention to resolve the challenges of overfitting, and varying input lengths of WSI analysis. HRDPath is a three-stream architecture integrating a DSMIL module, a global Vision Transformer (ViT), and a novel Local Window Transformer to extract multi-scale morphological representations (Extended Data Figure 1a). The model takes a set of patch-level features 𝐹 𝜖ℝ^𝑁×𝑑^ as input, where 𝑁 denotes the number of cleaned image patches from a slide and 𝑑 is the feature dimension. In the first stream, DSMIL identifies coarse-grained regions of interest (ROIs) and produces a slide-level feature embedding ℤ_𝐷𝑆𝑀𝐼𝐿_𝜖ℝ^𝑑^ through attention-based pooling. The ROIs are also used to guide local feature selection for the third stream. In the second stream, the full set of patch-level features is prepended with a CLS token and passed through a standard Vision Transformer. The final CLS token embedding ℤ_𝑉𝑖𝑇_𝜖ℝ^𝑑^ is extracted as a global semantic representation. The third-stream constructs 𝑘 fixed-size windows (of size 𝑤 × 𝑤) centered on the ROIs identified by DSMIL (Extended Data Figure 1d-f) where hyperparameter k is identified by experimentation. DSMIL’s lightweight architecture allowed faster training and quick identification of critical patch phenotypes which can serve as smooth additional supervision to explore by transformer encoders. Size of each window remains fixed to enable parallelism for efficiency. Each local window, after appending a CLS token and applying absolute positional encoding, has shape ℝ^(1+𝑤^^2)×𝑑^ and is processed independently through a lightweight transformer. The 𝑘 resulting CLS embeddings are averaged to yield the fine-grained local representation ℤ_𝑙𝑜𝑐𝑎𝑙_𝜖ℝ^𝑑^. The outputs from the three streams, ℤ_𝐷𝑆𝑀𝐼𝐿_, ℤ_𝑉𝑖𝑇_ , and ℤ_𝑙𝑜𝑐𝑎𝑙_ are concatenated to form a slide-level representation ℤ_𝑠𝑙𝑖𝑑𝑒_𝜖ℝ^3×𝑑^. For a patient with 𝑆 slides, these are concatenated across slides to yield a patient-specific tensor ℤ_𝑝𝑎𝑡𝑖𝑒𝑛𝑡_𝜖ℝ^𝑆×3×𝑑^ . A CLS token is then prepended, resulting in an input of shape ℝ^(1+𝑆×3)×𝑑^, which is passed through a patient-level transformer to model inter-slide dependencies and enhance cross-slide pathological pattern recognition. The output CLS token from this transformer (also shape ℝ^𝑑^) serves as the final patient-level embedding. This embedding is then passed to a multi-task prediction head for joint learning of two correlated downstream tasks: HRD status classification (binary), and HRD score regression (continuous). All model components are initialized using Kaiming uniform initialization with fixed random seeds and trained end-to-end using a unified optimization framework. Eight attention heads were utilized which was the number utilized in the original ViT paper^62^. The number of encoder layers of 2 was determined based on available computational resources and these are now included in our methods. Final HRDPath hyperparameters used for reported performance (transformer encoder depth: 2, attention heads: 8, dimension: 1024, k=8, w=32, dropout=0.1 (Uniform in all transformer encoder layers and multi-layer perceptron), learning rate: 1e-4, weight decay: 1e-5, batch size:1, number of training epochs: 150).

### Model Benchmarking Implementation Details

To ensure consistent and fair benchmarking, all models were evaluated using a standardized patient-level aggregation module. The input resolutions and adaptation strategies varied across models. All models except DeepHRD and ZoomMIL utilized features extracted at 15× magnification. The UNI model was evaluated exclusively at 15×, using a pretrained UNI2-h backbone as a frozen feature extractor to generate patch-level embeddings. All models will output the slide-level representation from their original architecture and aggregated by HRDPath’s patient-level transformer encoder for patient-level prediction. DeepHRD was implemented in a two-stage, non-end-to-end pipeline. Initially, DeepHRD was trained on 7.5× resolution patches to identify regions of interest (ROIs). The resulting model was then used to guide ROI selection for training a second DeepHRD model at 15× resolution. DeepHRD employs an ensemble of ResNet-18 backbones, and for each ensemble member, the top scoring patches were averaged to obtain slide-level features (following the original study). The final slide representation was derived by averaging across ensemble outputs. ZoomMIL also incorporates a multi-resolution approach: it first encodes WSIs at 7.5× magnification to identify ROIs, then re-encodes those regions at 15× resolution. The final slide-level representation consists of the concatenation of the 7.5× and 15× features (Originally ZoomMIL utilized three scales, including thumbnail, for fair comparison, it is fixed to two scales). This unified aggregation strategy ensures that model comparisons focus on differences in feature extraction and representation learning, rather than downstream fusion or classification methods.

### Intra-patient consistency analysis

Each patient in our dataset is associated with a variable number of slides. To assess the consistency and robustness of model predictions at the patient level, we designed a systematic experiment analysing how prediction accuracy varies with the number of slides available per patient. For each patient with *n* slides (where *n* ≥ 1), we evaluated model performance across all possible combinations of *i* slides (1 ≤ *i* ≤ *n*). For instance, for a patient with three slides labelled (a, b, c), when evaluating performance at *i* = 2, we considered all combinations of slide pairs: (a, b), (a, c), and (b, c). The model was independently run on each combination, and the resulting patient-level predictions were aggregated and compared to the ground truth to calculate average prediction accuracy.

We repeated this process across all patients for increasing values of *i*, ranging from 1 to the maximum number of slides observed in our dataset. The aggregated accuracy was computed by averaging performance across all patients with at least *i* slides.

To further investigate the impact of predictive confidence, we computed agreement scores for each prediction combination, defined as the proportion of constituent slide-level predictions. This approach allowed us to quantify not only the marginal benefit of incorporating more slides but also the extent to which consistency among slides correlates with improved prediction reliability. Importantly, inference is computationally efficient as each slide requires only ∼2.4 seconds to process.

### Morphology Analysis

We performed single-cell morphology analysis using Hover-NeXt pretrained on the PanNuke^63^ dataset. We qualitatively assessed several model variants and found that pannuke_convnextv2_tiny_3 version performs best. Cells were classified as neoplastic, inflammatory, or connective (stromal). Since Hover-NeXt was not originally trained on images of ovarian cancers, we validated its performance by comparing the model’s output to reference scores generated through manual qualitative assessment by a pathologist for 40 randomly selected WSIs. We observed good correlations for HoVer-NeXt-based scores and pathologist scores of neoplastic cell proportion (Spearman correlation = 0.67, p≤0.001) and stromal cell proportion (Spearman correlation = 0.76, p≤0.001).

### Morphology Feature Computation

Morphological and pathological patterns were captured by evaluating features including cell proportion and purity, neoplastic cell roundness, elongation, area, local cell density, and distance to the nearest immune cells (IND). Cell proportions were obtained by calculating the percentage of a certain cell type contained in each slide, while the purity of a certain cell type was the slide average of patch cell proportion. Cellularity is the total number of identified cells normalized to region area (patch or high-attention region). Roundness, elongation, and area were computed from segmented cell contours using the Python OpenCV library. The local cell distance (LCD) was calculated based on the inverse of the average Euclidean distance of cell centroids. The distance between neoplastic cells and immune cells (IND) was obtained by calculating the mean distance of the nearest 10 immune cells to each neoplastic cell. Z-score normalization was applied to the features, using the mean and standard deviation of the entire dataset. Outliers with an absolute z-score greater than 3 were filtered out. All features were then scaled to a range between 0 and 1. The morphology of neoplasms with different genomic alterations was computed across all anatomical locations (n=720 slides; HRD-: 350 slides; Other HRD+: 75 slides; *BRCA1/2*-mutated:197 slides; *BRCA*-methylated: 98 slides). To assess morphological features across different anatomical locations, 703 slides were analyzed, excluding those labeled ’other location’.

### VAttentionClusters Generation

HRDPath is a patient-level transformer-based deep learning model with two levels of attention: (i) patient-level attention to assign a score to each WSI to identify importance, and (ii) slide-level attention to demonstrate HRD-related regions within WSI. VAttentionClusters were defined based on HRDPath attention using a multi-stage approach (Fig 4e). DBScan clustering (epsilon=3.5) was applied based on the HRDPath attention score and spatial distribution to identify attention clusters within each WSI and filter patches with inconsistent spatial attention patterns. This resulted in the identification of 1542 attention clusters. To characterize semantically meaningful morphological patterns within high-attention regions, we implemented k-means clustering on identified attention clusters regarding their visual features. and qualitatively evaluated that it captures distinct phenotypes and tissue regions. This design enables a conceptual correspondence between clusters and attention heads, supporting interpretability under the hypothesis that each head captures distinct morphological signature ^64^.

### VAttentionCluster Enrichment Factor

The enrichment factor is the product of VAttentionCluster proportion and percentage of enrichment. The VAttentionCluster proportion is the ratio of the number of VAttentionClusters to all identified attention clusters in a slide. The percentage of enrichment is computed by dividing the regions of a specific VAttentionCluster occupied in a slide by the entire slide region.

All statistical analyses were performed using Python (v3.10) with SciPy, scikit-learn libraries. Model performance was evaluated using area under the receiver operating characteristic curve (AUROC), sensitivity and specificity. Associations between HRDPath attention and cellular morphological features extracted by HoVer-NeXt were evaluated using t-test with statistical significance defined as p≤0.05. Cellular morphological features are normalized and standardized, outliers (>±3 standard deviation) are dropped.

## Data and code availability

TCGA-OV and PTRC-HGSOC data is publicly available. The UWOV data is not publicly available due to patient privacy and consent restrictions but can be accessed upon reasonable request from the corresponding author.

## ACKNOWLEDGEMENTS

We acknowledge all members of the Sailem group. We acknowledge Iain McNeish for valuable early discussions. HS is funded by a Wellcome Career Development Award 225974/Z/22/Z. For the purpose of open access, the author has applied a CC BY public copyright license to any Author Accepted Manuscript version arising from this submission.

## AUTHOR CONTRIBUTIONS

Conceptualization, HS, CW, LS, KB; Methodology: CW, HS; Investigation: CW, HS, LS, KB, MD, MR, CNZ, RS, RA; Visualization: CW, HS; Funding acquisition: HS; Project administration: HS; Supervision: HS; Writing – original draft: CW, HS; Writing – review & editing: HS, CW, LS, KB

